# The Circadian Isoform Landscape of Mouse Livers

**DOI:** 10.1101/2024.10.16.618716

**Authors:** Alexander Zee, Dori Z.Q. Deng, Luciano DiTacchio, Christopher Vollmers

## Abstract

The mammalian circadian clock is an autoregulatory feedback process that is responsible for homeostasis in mouse livers. These circadian processes are well understood at the gene-level, however, not well understood at the isoform-level.

To investigate circadian oscillations at the isoform-level, we used the nanopore-based R2C2 method to create over 78 million highly-accurate, full-length cDNA reads for 12 RNA samples extracted from mouse livers collected at 2 hour intervals. To generate a circadian mouse liver isoform-level transcriptome, we processed these reads using the Mandalorion tool which identified and quantified 58,612 isoforms, 1806 of which showed circadian oscillations. We performed detailed analysis on the circadian oscillation of these isoforms, their coding sequences, and transcription start sites and compiled easy-to-access resources for other researchers.

This study and its results add a new layer of detail to the quantitative analysis of transcriptomes.

## INTRODUCTION

The mammalian circadian clock is an autoregulatory feedback process that is responsible for 24 hour oscillations in mRNA and protein levels. Circadian clock genes are driven by the transcription factors BMAL1 and CLOCK (*1–3*). BMAL1 and CLOCK heterodimerize to form large multiprotein complexes. This complex drives the transcription of *Cry,* and *Per*, and in turn, their protein products inhibit BMAL1/CLOCK transcriptional activity thus forming a negative feedback loop (*4–9*).

In mammals, the circadian system is composed of a collection of cell- and tissue-level oscillators whose phase is coordinated by a combination of light and non-light stimuli (*10*). (*11*, *12*).

In the mouse liver, the circadian clock and feeding/fasting cycles together generate robust oscillations in thousands of genes (*13–15*), including those encoding proteins involved in every major metabolic pathway. Indeed, the molecular mechanisms that integrate the molecular clock with the genomic circuitry responsible for large-scale oscillations include transcription factors that are master regulators of energy metabolism, including CREB1, and SREBP1 (SREBF1). Consequently, circadian dysregulation due to environmental or genetic causes is an important factor in the development of metabolic diseases (*16–20*).

Many studies have investigated genome-wide transcriptional oscillations in mouse liver (*10*, *21–23*). However, due to technical limitations, these studies have been limited to the level of genes or individual elements like splice sites and junctions. Moving beyond the analysis of genes and their individual elements and towards the analysis of transcript isoforms has the potential to increase our understanding of circadian oscillations.

In addition to revealing how splice-junctions are chained to form the isoforms themselves, transcript isoforms contain information on 1) their complete coding sequences (CDSs) and 2) the location of their transcription start sites (TSSs). While CDSs reveal the actual protein isoforms causing downstream effects, TSSs pinpoint the sites of transcriptional regulation.

Advancements in long-read sequencing have now enabled the accurate and high-throughput sequencing of full-length cDNA (*24*, *25*). Further, advancements in isoform identification pipelines allow for the identification and quantification of isoforms, including TSSs (*26–28*). Finally, advancements in tools for functional isoform classification and annotation allow for the identification of CDSs in isoforms (*29*).

Here, we took advantage of these advancements and investigated circadian oscillation in expression on the isoform-level in the mouse liver. Using R2C2, a long read sequencing method (*24*, *30–34*), which increases accuracy and reduces length bias of ONT sequencers, we generated over 78 million accurate, full-length cDNA (fl-cDNA) reads from mouse livers across 12 timepoints spaced at 2 hours. We analyzed these fl-cDNA reads using the Mandalorion isoform identification pipeline (*35*) to identify and quantify isoforms and their TSSs, then used SQANTI3/IsoAnnotLite (*29*) to functionally annotate them and add CDSs. Further, we detected the circadian oscillation patterns of isoforms, CDSs, and TSSs using MetaCycle (JTK) (*36*), characterized these oscillations, and generated resources to facilitate the exploration of this dataset.

## RESULTS

### Generating Highly Accurate Transcriptome Data Across a Circadian Time Course

To generate a circadian time course of mouse liver RNA samples for fl-cDNA sequencing, we started by keeping male C57BL/6 mice in standard 12 hr Light/Dark cycles. We then transitioned these mice to constant darkness and, beginning at CT18, sacrificed pairs of mice at 2 hr intervals over 24 hours to collect their livers (Fig. 1).

**Fig. 1:**
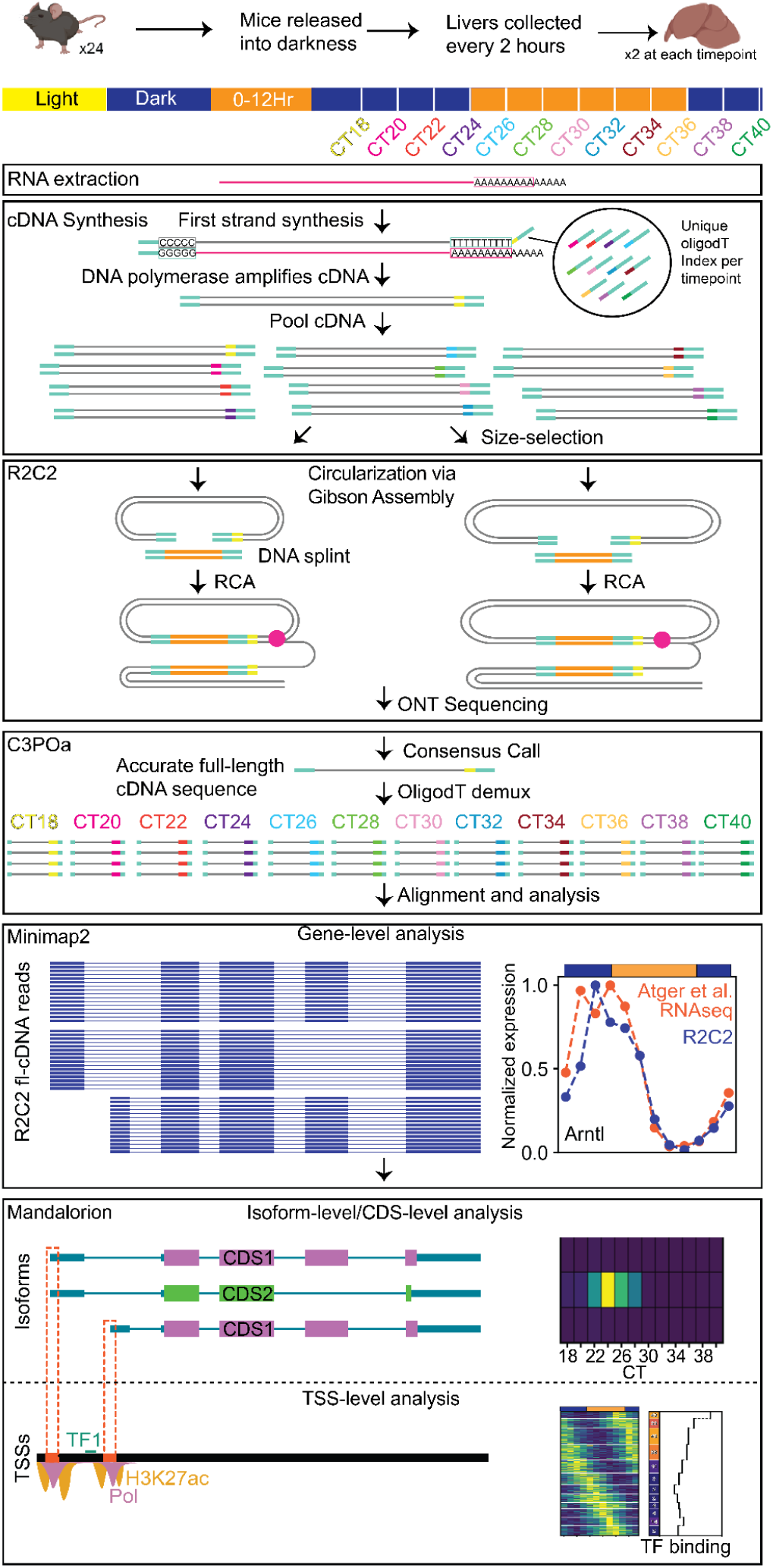
Study Overview. Full-length cDNA was created from RNA extracted from mouse livers at 2 timepoints. Pooled cDNA was prepared for ONT sequencing using the R2C2 method. ONT raw reads were consensus called using C3POa, and then demultiplexed based on Oligo(dT) index sequence. Demultiplexed reads were then aligned and used to investigate gene-, isoform-, CDS- and TSS-level circadian oscillations using several computational tools including Mandalorion.

After extracting RNA from the two liver samples from each timepoint, we constructed fl-cDNA libraries from each RNA sample. We prepared fl-cDNA using indexed oligo(dT) primers during first strand synthesis, with a unique index being used for each RNA sample. This allowed us to pool fl-cDNA from all timepoints after PCR amplification to minimize batch effects from downstream processing (Fig. 1).

Some of the fl-cDNA pool was size-selected for molecules >2.5kb in length by gel electrophoresis to increase sequencing coverage of longer transcripts. We then used the R2C2 method to prepare size-selected and non-size-selected samples for ONT sequencing (Fig. 1).

R2C2 uses a DNA splint to circularize the cDNA with a version of Gibson assembly - NEBuilder HIFI DNA Assembly. The circularized cDNA is then amplified using phi29 and results in a linear dsDNA with multiple copies of the same cDNA sequence, separated by the DNA splint (Fig. 1).

We sequenced this concatemeric DNA on MinION and PromethION sequencers using R9.4/SQK-LSK110 or R10.4/SQK-LSK114 pore chemistry/library preparation kits. We basecalled the raw signal using Guppy (v6) (*37*) and used the C3POa (v3.2) tool to process the resulting raw reads. C3POa separates the concatemeric raw reads into subreads and uses abpoa (*38*), minimap2(*39*), and racon(*40*) to combine these subreads to produce consensus reads. C3POa also demultiplexes consensus reads into their respective timepoints based on their oligo(DT) index sequence (Fig. 1). In total, we generated 78.7 million highly-accurate full length cDNA reads with an average of 6.55 million reads per time point. Across all timepoints, the median accuracy, i.e. identity to the mm39 genome, of these reads was 99.8% and their length was ∼1.5kb. Due to the indexing and pooling strategy, data was consistent and well distributed across the 12 different time points (Fig. S1).

Overall, this showed that our experimental setup generated an evenly sequenced, highly accurate fl-cDNA data set for our circadian time course which we subsequently used for gene-, isofom-, CDS- and TSS-level analysis (Fig. 1).

### Detecting Gene-Level Circadian Oscillations

To validate that our experimental setup also generated data quantitative enough to detect circadian oscillations, we first performed gene-level analysis. To quantify gene expression we aligned the R2C2 reads to the mm39 version of the mouse genome using minimap2. Separately for each timepoint, we then used featureCounts (*41*) to count the number of R2C2 reads that were aligned to each gene present in the GENCODE vM30 annotation. Finally, we normalized these read counts using quantile normalization with NormSeq (*42*).

We then examined the gene expression of well known oscillating genes in this normalized data set: *Arntl (Bmal1), Dbp, Insig2, Nfil3, Nr1d1, Rorc, Elovl3, Rgs16,* and *Cry1* (Figure 2A). We found that their expression levels indeed oscillated in our data set and that their oscillation patterns (*43–47*) and peaks matched those of a publicly available short-read RNA-seq liver time course (Atger et al.) (*48*).

**Fig. 2:**
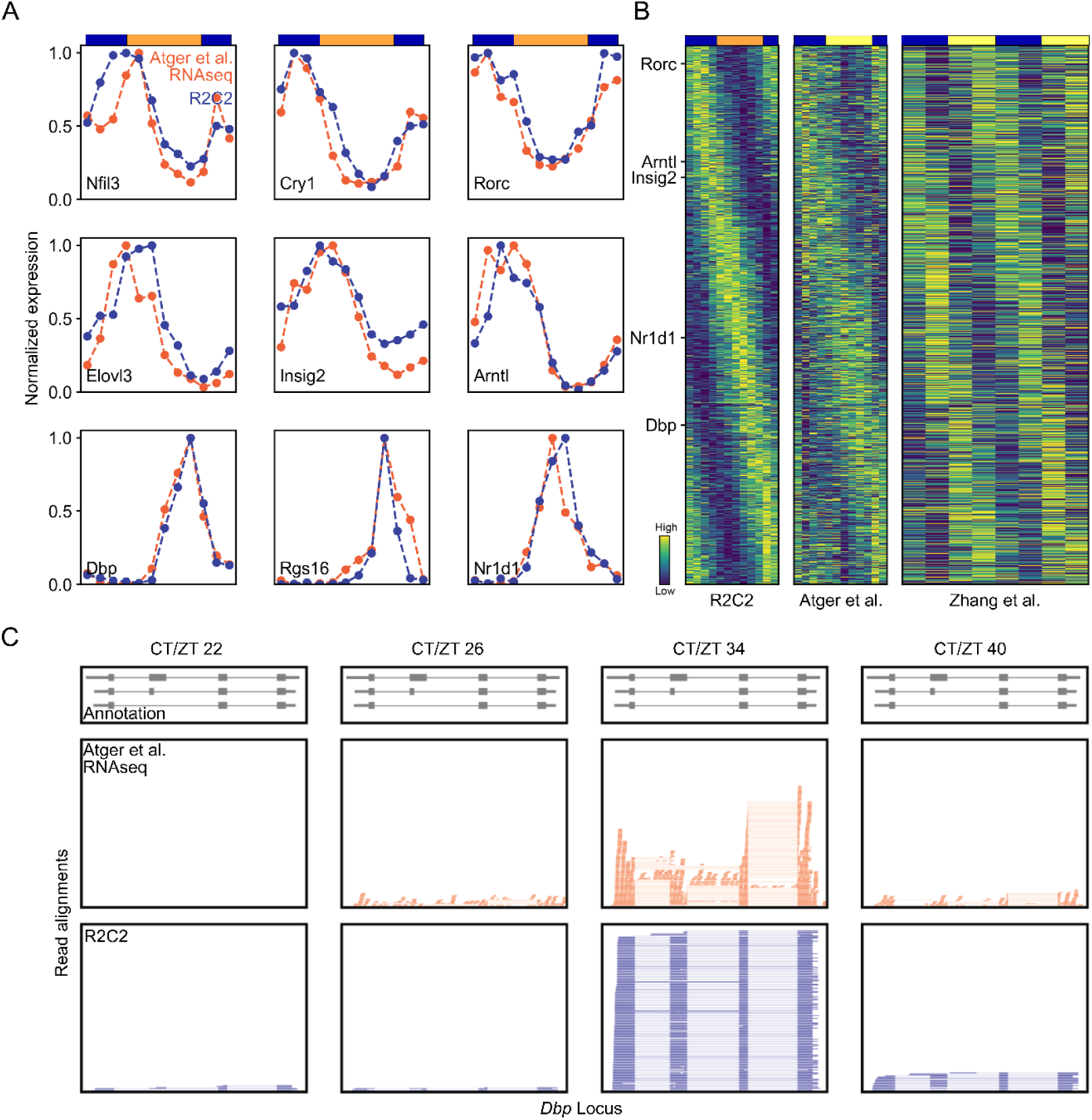
Detecting Gene-Level Oscillations Using R2C2. A) Gene level oscillation analysis of known circadian genes. Normalized gene expression is shown for both R2C2 fl-cDNA and the Atger et al. RNA-seq data set. Circadian time (CT) is indicated on top as blue and orange rectangles with the blue to orange transition indicating CT24 and orange to blue transition indicating CT36. B) Heatmaps showing the gene expression of the 1068 genes scored as oscillating in the R2C2 data set. Genes were sorted by the peak phase in the R2C2 data set. In addition to the R2C2 data set, gene expression is shown for the Atger et al. and Zhang et al. data sets. Time is indicated as in A) with yellow rectangles indicating Zeitgeber time (ZT) instead of Circadian time (CT). C) GenomeBrowser style visualizations of read alignments at the *Dbp*. Top panel: Basic GENCODE vM30 annotation. Middle panel: RNA-seq read alignment from the Atger et al. data set. Bottom panel: fl-cDNA R2C2 consensus read alignments.

To perform systematic, genome-wide analysis, we used metaCycle(*36*) with JTK_CYCLE(*49*) to evaluate oscillation across all genes. By applying an adjusted p-value cutoff of 0.05, we found 1068 genes with oscillating expression levels (Figure 2B). Visualization of these oscillating genes showed that their peak phases were distributed across the entire 24 hours of the time course. We also found that the oscillations of these genes in the R2C2 data set are similar to their oscillations in two publicly available short-read RNA-seq datasets (Atger et al. and Zhang et al.) (Fig. 2B) (*22*, *48*).

We then compared the list of genes scored as oscillating in the R2C2 data set to the lists of genes scored as oscillating in the two publicly available short-read RNA-seq data sets. After we reanalyzed these two data sets with the same featureCounts to NormSeq to MetaCycle (JTK) workflow, we found that 435 of 1068 oscillating genes in the R2C2 data set were also oscillating in at least one of the other two data sets. Importantly, this compares well to comparison between the two publicly available short-read RNA-seq data sets. Of the 1739 and 1058 genes with oscillating gene expression in Atger et al. and Zhang et al. short-read data sets, respectively, only 213 oscillate in both data sets.

Overall, this showed that the data generation and quantitation approach we designed and employed is quantitative enough to enable robust detection of oscillating gene expression. It shows core clock genes oscillating with the expected phases and the oscillating genes it detects overlap well with publicly available short-read RNA-seq data sets.

### Identifying and isoforms, CDSs, and TSSs

Importantly, however, R2C2 is not limited to gene-level analysis. In contrast to RNA-seq reads, R2C2 reads span entire transcripts. R2C2 and RNA-seq reads aligned to the *Dbp* locus highlight this difference (Fig. 2C). Because of that, R2C2 data enables the identification and quantification of transcript isoforms, including coding sequences (CDSs) and transcript start sites (TSSs).

To move beyond gene-level analysis of circadian oscillation of expression, we processed the R2C2 fl-cDNA data set with Mandalorion (v4.5). Mandalorion has shown excellent isoform identification performance in the LRGASP consortium (*28*) comparison and has since been expanded to quantify isoforms as well (*35*). The newest version of Mandalorion (v4.5) also enables the quantification of transcription start site (TSSs) usage and includes powerful tools for isoform visualization.

By processing the entire ∼78 million read data set, Mandalorion identified 58,612 isoforms and quantified them across timepoints. 47,111 of these isoforms were predicted to be protein-coding and contained 29,154 unique CDSs. Further, these isoforms used 24,645 non-overlapping transcription start sites (TSSs).

### Detecting isoform oscillations

First, we investigated oscillating isoform usage and how it can increase the resolution of the circadian transcriptome. Similar to gene-level analysis, we normalized isoform expression time course using NormSeq and processed the normalized data using MetaCycle (JTK). Using an adjusted p-value cutoff of 0.05, we identified 1806 isoforms with oscillating expression (Fig. S2). These 1,806 isoforms were expressed by 1,415 genes, 553 (39%) of which were oscillating at the gene level.

To investigate this surprisingly low level of overlap, we calculated the ratio of expression of oscillating isoforms/all isoforms for genes expressing at least one oscillating isoform. This analysis showed a bimodal distribution with the majority of genes showing a ratio of either close to 0 or 1 (Fig. 3A left). The higher this ratio was, the more likely the gene itself was to be scored as oscillating (Fig. 3A right).

**Fig. 3.**
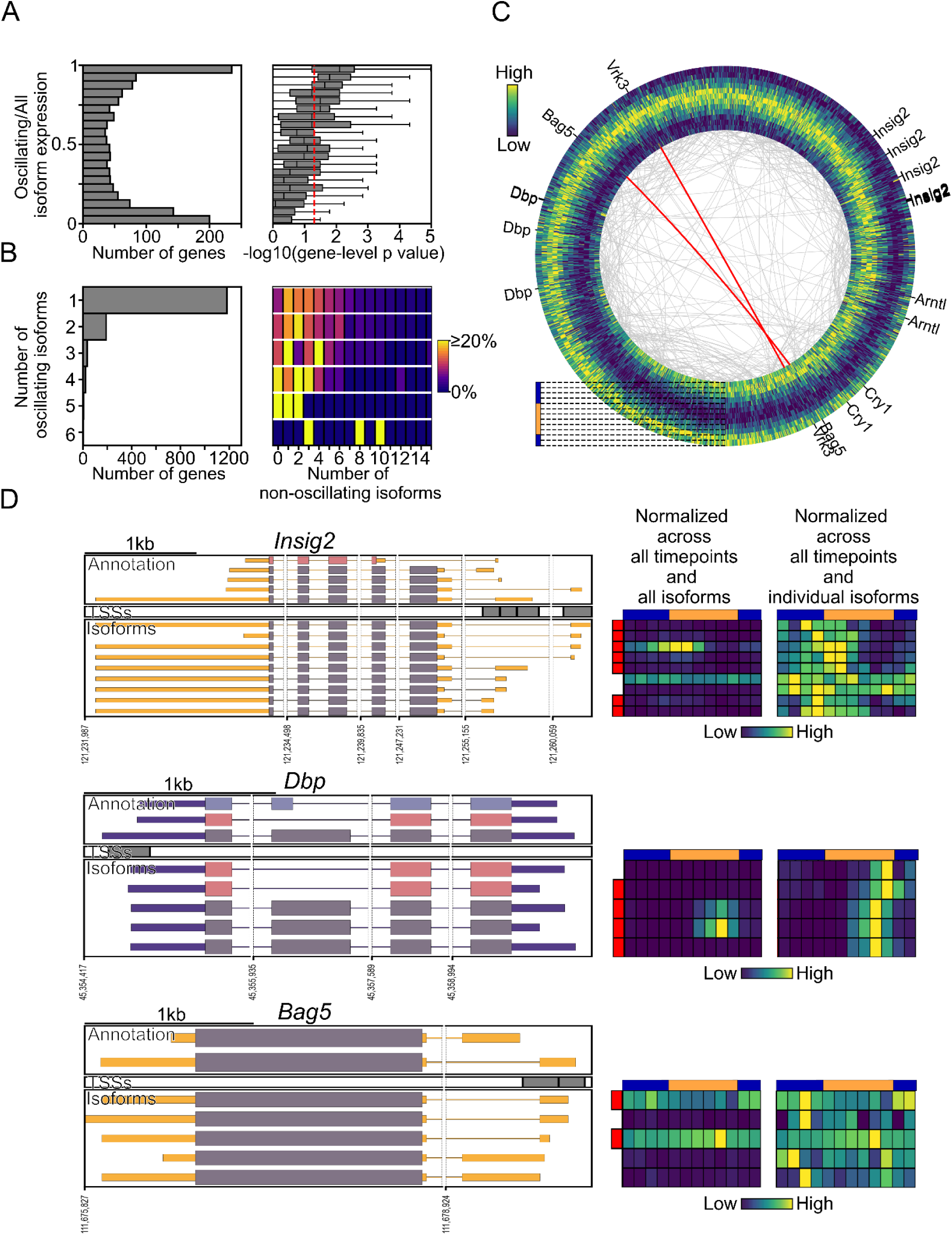
Detecting isoform-level oscillations. A) Left: Histogram of genes with at least one oscillating isoform binned by their expression ratio of oscillating isoforms to all isoforms. Right: Boxplots showing gene-level p-Value (-log10 converted) distributions of across the corresponding bins on the left. The dashed red line indicates the p-value cutoff of 0.05 B) Left: Genes with at least one oscillating isoform are grouped based on the number of oscillating isoforms they express. Right: Heatmaps showing the number of non-oscillating isoforms expressed by the corresponding groups on the left. C) Circular heatmap showing the 1806 oscillating isoforms. CT18 is on the inside, CT40 on the outside. Isoforms of select genes are indicated on the outside. Isoforms of the same gene are connected by grey lines on the inside with the exception of isoforms of *Vrk3* and *Bag5* which are connected by red lines. D) Left: Genome Browser style visualizations of *Insig2*,*Dbp*, and *Bag5* featuring GENCODE annotations, TSS and isoforms. The same CDSs are displayed in the same color. Right: Heatmaps of isoform expression normalized across all timepoints and either all isoforms or within each isoform. Oscillating isoforms are marked by red rectangles.

Further analysis showed that the majority of genes expressing oscillating isoforms expressed a single oscillating isoform but often several non-oscillating isoforms (Fig 3B). However, 254 genes expressed more than one oscillating isoform. In the majority of cases, isoforms within each gene showed similar expression patterns as seen for multiple isoforms of *Arntl*, *Cry1*, and *Dbp* (Fig. 3C,D). There were only two genes - *Vrk3*, *Bag5 (Fig. 3C,D) -* that contained oscillating isoforms 1) whose peak-phases were more than 8 hours apart 2) that had similar overall expression levels (<2 fold difference) 3) and made up the majority of their gene’s expression (>75%).

Overall, this suggested three things. First, that many genes expressed a mix of oscillating and non-oscillating isoforms which in turn lowers the amplitude of gene-level oscillations and thereby increases their gene-level Metacycle adjusted p-values. *Insig2* is an example of this, expressing two dominant isoforms, one oscillating and one non-oscillating (Fig. 3D top). Second, that genes expressing more than one oscillating isoform are likely to have similar though not necessarily identical expression patterns. *Dbp* is an example of this with a less expressed oscillating isoform, peaking two hours after the most expressed oscillating isoform (Fig. 3D middle). Third, that genes expressing two equally abundant oscillating isoforms that show opposing peak-phases are very rare. *Bag5* is one of only 2 genes in our data set that show this behavior with two isoforms peaking ∼11 hours apart. Interestingly, these isoforms contained the same CDS but used different TSSs (Fig 3D bottom).

### Identifying Oscillating CDSs

Because the majority of isoforms and especially oscillating isoforms were classified as protein coding, we next performed an analysis on the level of coding sequences (CDSs). For this, we used the coding classification and CDS as determined by SQANTI3 for each isoform. The 47,111 isoforms classified to be protein-coding contained 29,154 unique CDSs.

To evaluate CDS expression, we combined read counts of all isoforms containing the same CDS. To see whether these combined expression values aligned with CDS characteristics, we stratified CDSs based on whether they were present in GENCODE and whether they were scored as nonsense mediated decay (NMD) targets by SQANTI3.

The 12,814 CDSs that were present in the GENCODE annotation and were more highly expressed than the 16,340 CDSs that weren’t (Fig. 4A). This is consistent with more highly expressed isoforms and their CDSs being more likely to be previously annotated. Conversely, the 1,594 CDSs which were scored as likely NMD targets were expressed lower than the ones that weren’t (Fig. 4A). This is consistent with NMD targets being actively degraded and therefore present at a lower rate.

**Fig. 4.**
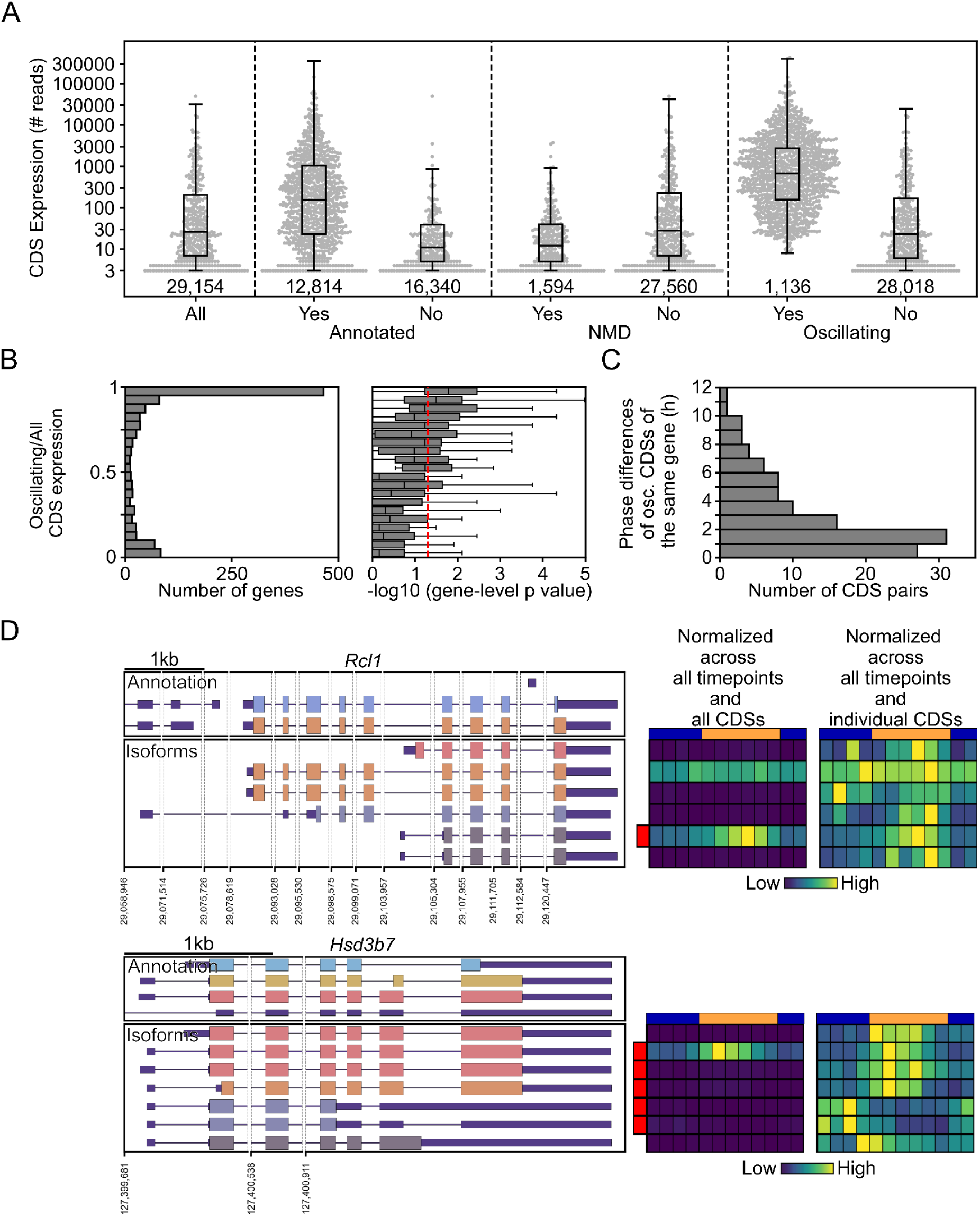
Detecting CDS-level oscillations. A) Swarmplots and overlaid boxplots showing expression levels of all CDSs and CDSs categorized by whether they are 1) present or not-present in the GENCODE annotation, 2) predicted NMD targets or not, and 3) scored as oscillated or non-oscillating. Numbers below each swarmplot indicate the total number of CDSs in each group. B) Left: Histogram of genes with at least one oscillating isoform binned by their expression ratio of oscillating isoforms to all isoforms. Right: Boxplots showing gene-level p-Value (-log10 converted) distributions of across the corresponding bins on the left. The dashed red line indicates the p-value cutoff of 0.05 C) Histogram showing pairs of CDSs expressed by the same gene binned by the difference in their peak phase. D) Left: Genome Browser style visualizations of *Insig2*, *Dbp*, and *Bag5* featuring GENCODE annotations, TSS and isoforms. The same CDSs are displayed in the same color. Right: Heatmaps of isoform expression normalized across all timepoints and either all isoforms or within each isoform. Oscillating isoforms are marked by red rectangles.

Next, to identify oscillating CDSs, we used NormSeq to normalize CDS read counts and MetaCycle (JTK). Using an adjusted p-value cutoff 0.05 we found 1,136 oscillating CDSs. These 1,136 oscillating CDSs were generally very highly expressed (Fig. 4A) and belonged to 1036 genes, 506 of which were determined to be oscillating at the gene-level. Similar to our isoform-level analysis, we investigated why there were genes expressing oscillating CDSs that weren’t scored as oscillating themselves. To do this, we quantified the ratio of oscillating/all CDS in genes with at least one oscillating CDS. We again found a bimodal distribution, but the majority of genes showed an oscillating/all CDS ratio close to 1. (Fig. 4B left). As with isoform-level analysis, the higher this ratio was, the more likely the gene itself was to be scored as oscillating (Fig. 4B right). Finally, if a gene expressed more than one oscillating CDS, they were very likely to have similar, yet not necessarily identical peak phases (Fig. 4C).

In summary, CDSs behaved similar, yet not identical to isoforms in our analysis. Considering CDS expression was compiled from isoform expression, this was expected. We identified genes like *Rcl1* that expressed both oscillating and non-oscillating CDSs at similar levels (Fig. 4D top). The *Hsd3b7* gene on the other hand expressed three oscillating CDSs. One major and one minor CDS with similar peak phase as well as a minor CDS which peaked 6 hours earlier and was predicted to be an NMD target (Fig. 4D bottom).

These two examples show that CDS-level analysis can add valuable detail to the analysis of oscillating gene expression and should be of particular interest to scientists with specific genes of interest. We have therefore generated Genome Browser style images (as in Fig. 3D and 4D) for all expressed genes in our data set and are supplying them as supplemental data.

### Identifying Oscillating Transcription Start Sites

Next, we investigated oscillating TSS usage and how it can uncover how oscillating transcription is regulated. To do so, we performed a detailed analysis on 24,645 TSSs identified by Mandalorion.

We systematically validated these 24,645 TSSs using refTSS (*50*) data as well as publicly available circadian ChIP-seq time course data for H3K4me3(*51*), H3K27ac, and PolII(*52*) - all associated with promoters and TSSs (*21*, *53*). For refTSS, we used the provided BED file to calculate the overlap of refTSS peaks with the TSSs we identified. For ChIP-seq, we aligned the reads using BWA (*54*) and calculated the normalized density of the three marks around the TSSs we identified.

We found that refTSS peaks were highly correlated with the TSSs - ∼50% of TSSs overlapped with a refTSS peak (Fig. 5A top). Further we found that histone marks H3K4me3 and H3K27ac showed the asymmetric, stereotypical density around our TSSs expected for sites of active transcription initiation: A smaller increase in density upstream of the TSSs was followed by a drop directly over the TSS - likely due to an absence of histones - in turn followed by a larger increase in density just downstream of the TSSs (Fig. 5A top). Finally, PolII signals showed high density over the TSSs, peaking approximately 60nt downstream of the TSSs (Fig. 5A top). We visualized and validated these ChIP-seq enrichment patterns by plotting normalized densities for the 10,000 most expressed TSSs (Figure 5A bottom). Together, this showed that the location of the TSSs Mandalorion had identified were in concordance with independent TSSs determining assays (refTSS) and epigenetic markers for active promoters (H3K4me3, H3K27ac, PolII).

**Fig. 5.**
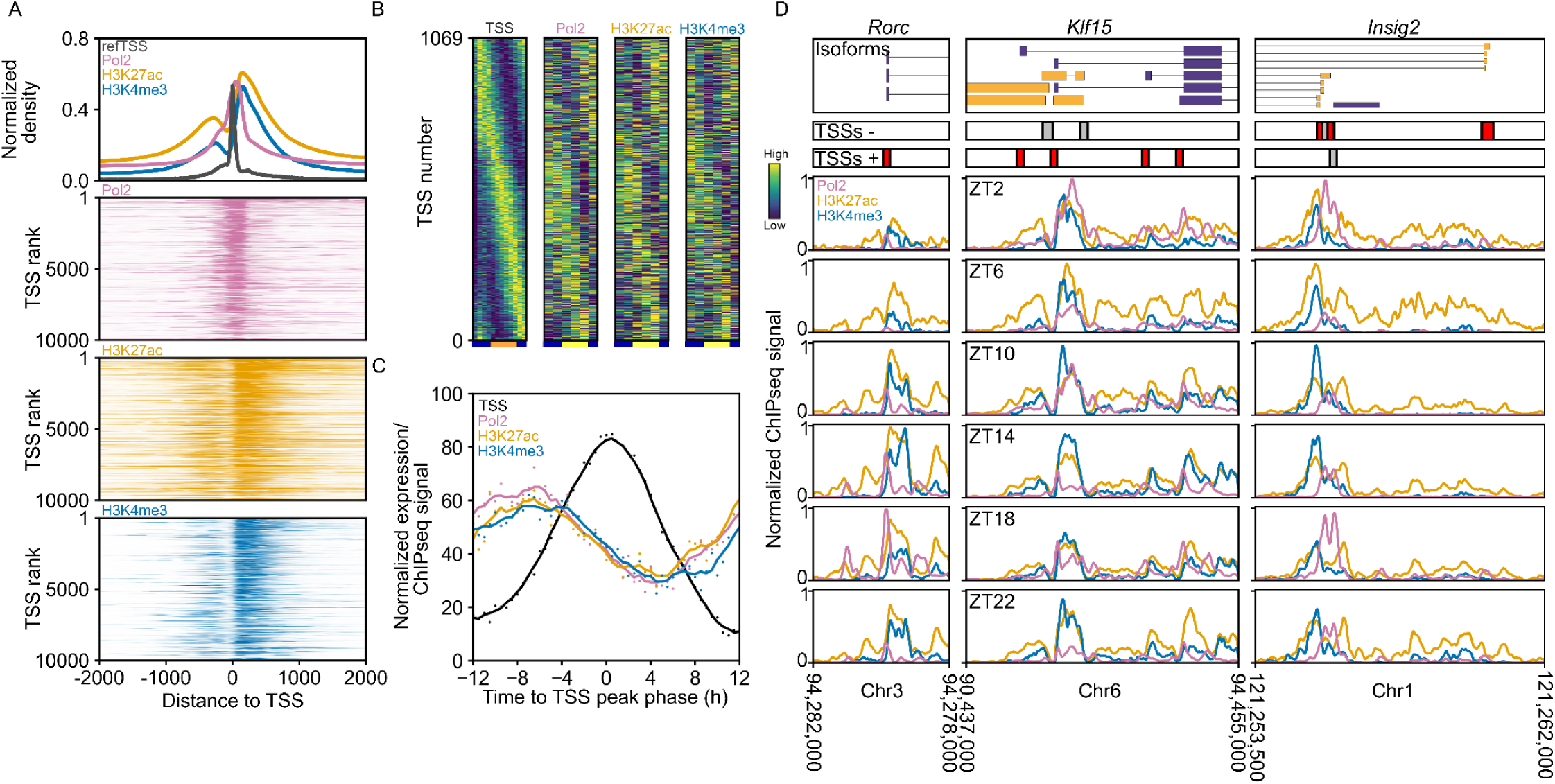
Identifying TSSs and validating their oscillations. A) refTSS peak density and normalized ChIP-Seq read density for PolII, H3K4me3, and H3K27ac is shown around identified TSS. On top, density data is shown in aggregate. Below, normalized ChIP-seq density is shown as heatmaps for the 10,000 highest expressed TSS sorted in descending order. B) Heatmaps showing the expression of the oscillating 1069 TSS sorted by the peak phase in the R2C2 data set. Normalized PolII, H3K4me3, and H3K27ac ChIP-seq density is shown for the same TSSs. Time is indicated as in Fig. 3B. C) Peak-phase adjusted and normalized TSS expression and PolII, H3K4me3, and H3K27ac ChIP-seq are shown as line graphs. D) Top, the 5’ ends of isoforms for Rorc and Insig2 are shown, below, TSSs as called by Mandalorion are shown as boxes and colored red if they were scored as oscillating. At the bottom, normalized ChIP-seq read density for 6 time points is shown.

Next, we wanted to detect and validate oscillating TSSs. To detect oscillating TSSs, we followed the same workflow as we did for gene and isoform oscillations. We used NormSeq to normalize TSS read counts reported by Mandalorion and then used MetaCycle (JTK) to detect significant TSS oscillations. Using an adjusted p-value cutoff 0.05 we found 1069 oscillating TSSs (Figure 5B).

These 1069 oscillating TSSs belonged to 988 genes, 580 of which were determined to be oscillating at the gene level. This 54% oscillating TSS/oscillating gene overlap was higher than the 39% oscillating isoform/oscillating gene overlap. Consistent with that and similar to what we observed with CDSs, most genes that contained at least one oscillating TSS showed an oscillating TSS/all TSS ratio close to 1 (Fig. S3).

To further validate TSSs oscillations, we visualized them using heatmaps. We then also created heatmaps for H3K4me3, H3K27ac, and PolII for the genome regions around these same TSSs. This showed that TSSs oscillated with peak phases throughout the day and that H3K4me3, H3K27ac, and PolII mirrored these oscillations, albeit more noisily (Fig. 5B). Once averaged across all oscillating TSSs and phase-adjusted, H3K4me3, H3K27ac, and PolII peaks preceded TSS peaks by 2-4 hours which is in line with previous studies performed using RNA-seq at the gene level (Fig. 5C).

Together, these results indicate that we can accurately pinpoint the location of TSSs and detect their oscillations.

Three examples highlight how fl-cDNA can add detail to ChIP-seq data for identifying and quantifying TSS oscillations: First, at the *Rorc* locus, Mandalorion identified one TSS shared between 3 isoforms. The TSS as well as all 3 isoforms were determined to be oscillating. We also see a strong oscillation of H3K4me3, H3K27ac, and PolII levels in the promoter region associated with this TSS (Figure 5A). Second, at the *Klf15* locus, Mandalorion identified 4 oscillating TSSs with phases several hours apart. However, because these TSSs are closely spaced, the resolution of ChIP-seq fails at clearly delineating these areas. Third, at the *Insig2* locus, Mandalorion identified 4 different TSSs shared between its 9 isoforms. Three of these TSSs were determined to be oscillating, as were the isoforms associated with them. Again, we see oscillations in H3K4me3, H3K27ac, and PolII in the promoter regions. However, the resolution of the ChIP-seq assay is again not sufficient to locate and differentiate oscillating and non-oscillating TSSs.

This shows that while oscillating ChIP-seq of epigenetic markers and PolII can help validate TSSs location and oscillation, TSSs-level expression data is more precise at locating and quantifying TSSs and oscillations in their usage.

### Detecting circadian oscillations in Transcription Factor activity at TSSs

Having established the ability to determine the location and usage of TSSs, we next aimed to establish the relationship between transcription factors (TFs) and oscillating TSSs to quantify circadian oscillation in the activity of these TFs.

First, we used k-means (*55*) clustering to group similarly oscillating TSSs into 12 clusters. Then we downloaded publicly available TF ChIP-Seq data, which contained transcription factor occupancy peaks - i.e. binding sites - for mouse liver, from ChIP-Atlas.org (*56*). Next, we determined what percentage of 1) all 24,645 TSSs and 2) the TSSs in each of the 12 clusters overlapped with the binding sites of each TF ChIP-seq experiment. We used these percentages to identify which TF ChIP-seq experiments were at least 3-fold enriched in one of the 12 TSSs clusters when compared to all 24,645 TSSs. Then we used these enriched TF ChIP-seq experiments to calculate the average percentage of TSSs in each cluster overlapping with TF binding sites (Fig. 6).

**Figure 6.**
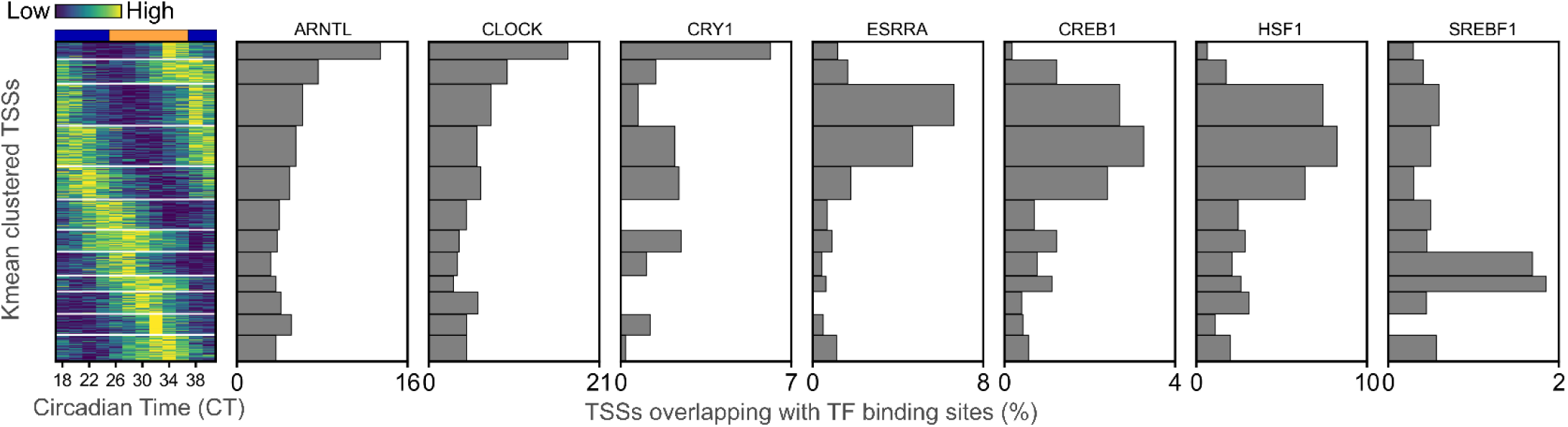
Quantifying circadian oscillations in transcription factor activity. Left) Heatmap of oscillating transcription start sites, clustered using K-means. Right) Core clock and metabolic TF enrichment across the 12 clusters: On top, the name of the TF and to its left, the total number of ChIP-seq experiments whose binding sites were significantly enriched in any of the 12 clusters are shown. Below the total number, the number of ChIP-seq experiments whose binding sites were significantly enriched in each of the 12 clusters is shown. The line graph on the right shows the average percentage of TSSs in each cluster that overlap with a binding site of any of those ChIP-seq experiments.

In this way, we found that ARNTL/BMAL1 binding sites were most likely to overlap with oscillating TSSs peaking around CT36. Indeed, ARNTL has previously shown to bind promoters around CT32-36 (*57*). The core clock TFs CLOCK and CRY1 produced highly similar patterns. CLOCK, of course, is known to dimerize with ARNTL to form an transcriptional activator while CRY1 is known to bind and repress that ARNTL/CLOCK dimer.

Next we looked at TFs known to control metabolic processes.

ESRRA, CREB1,and HSF1 are known to be activated during periods of fasting and to show circadian oscillation in their activity due to the feeding rhythms. Indeed, binding sites of these TFs were most likely to overlap with oscillating TSSs that started increasing at CT34 when mice were fasting and peaked at CT38 at which point mice have been feeding.

SREBF1, on the other hand, is known to be activated by feeding. Consistent with that, we found that binding sites for SREBF1 were most likely to overlap with TSSs that started increasing at CT20 after mice had fed and peaked around CT28.

Overall, this analysis shows that we can use our fl-cDNA data set to determine the exact location of TSSs and precisely quantify their usage. Coupled with public transcription factor occupancy data, we can then map when TFs are active and which TSSs they are involved in regulating.

## DISCUSSION

Over the last two decades, circadian oscillations in transcription have been investigated with ever improving methods - from RT-qPCR to Microarrays to RNA-seq. Yet, none of these methods has been capable of identifying and quantifying isoforms. This has left a large gap in our understanding of the circadian transcriptome.

Here, we used nanopore-based R2C2 full-length cDNA sequencing to fill that gap in the mouse liver.

We used the R2C2 to generate over 78 million fl-cDNA reads and the Mandalorion isoform identification tool to analyze these reads and produce a highly detailed isoform-level transcriptome. Our analysis identified 58,612 isoforms which contained 29,154 unique coding sequences (CDSs) and used 24,645 non-overlapping transcription start sites (TSSs). The high number of fl-cDNA reads we generated meant this isoform-level transcriptome contained a large number of new, unannotated elements. Indeed, 28,456 (49%) isoforms, 16,340 (56%) CDSs, and 10,917 (44%) TSSs, were not present in the GENCODE annotation. Further, many of the new elements were lowly expressed (<1RPM) (Fig. S4) indicating they would have been missed with lower overall read counts.

The design of our sequencing approach generated highly quantitative data which in turn allowed us to identify 1,806 isoforms, 1,136 CDSs, and 1,069 TSSs that showed circadian oscillations in their expression/usage. The isoform-level analysis showed that oscillating isoforms are often accompanied by non-oscillating isoforms expressed by the same gene - this, in turn, blunts gene-level oscillations. The CDS-level analysis combined isoforms based on the CDSs they contained, i.e. the protein they encoded. Beyond characterizing and cataloging CDS oscillations, we believe this analysis provides value to researchers which will now be able to perform more targeted studies. Finally, the TSS-level analysis provided insight into the regulation of circadian oscillations by enabling us to use public ChIP-seq data to identify the circadian activity of transcription factors (TFs). This is notable because this approach did not rely on timed collection of ChIP-seq data and could therefore be performed using ChIP-seq data generated from just one or pooled samples. This would simplify TF ChIP-seq experiments while still allowing the characterization of TF dynamics.

There are still inherent technical limitations of long-read sequencing and cDNA preparation that future studies will need to overcome.

First, while 6 million reads on average per time point was sufficient to capture robust circadian oscillations in medium to highly expressed isoforms, future studies should target 5 to 10 times as many reads to achieve parity with the approximate read numbers used in short-read RNA-seq studies and capture lowly expressed isoforms. The consistently increasing number of reads produced by ONT and PacBio sequencers is bound to make this feasible.

Second, while the average read length we achieved with R2C2 in this study are far longer than standard ONT cDNA sequencing (*58*) and on par with PacBio Iso-Seq and Kinnex, they still fail to capture the longest of mouse transcript isoforms. A notable example of this are isoforms of the *Per1, Per2, and Per3* genes, which at ∼6kb were too long to be captured in this study. Future studies could potentially use improved reverse transcriptases (*59*) and polymerases, as well as more aggressive size-selection to address this.

Despite these limitations, this data set provides an unprecedented look at circadian oscillations and their regulation in mouse liver. To make this data set and our accompanying analysis as easily accessible for researchers as possible, we provide detailed supplemental data files, genome annotations in the form of gtf files, as well as Genome Browser-like images as in Fig. 3 and 4 for all isoforms/genes expressed in our data set. We hope that researchers will use these resources to gain more detailed insights into circadian oscillations in their gene(s) of interest.

In summary, by generating and analyzing highly-accurate and quantitative full-length cDNA sequences we increased the resolution of circadian genomics analysis. The isoforms, CDSs, and TSSs we identified and quantified add new layers of detail to the circadian oscillations of gene expression. Finally, the fl-cDNA/R2C2/Mandalorion approach we employed in this study can be applied to other tissues or organisms with few or no modifications and even small tissues like the SCN should easily provide enough RNA for the Smart-seq2(*60*)-based fl-cDNA generation approach. This study therefore serves as a blueprint for future studies into circadian oscillations at the isoform-level.

## METHODS

### cDNA synthesis

RNA was extracted from flash frozen mouse livers using TRIzol. Approximately 200ng of total RNA from each sample was then mixed with dNTPs, a template switch oligo (TSO), and an oligo(dT) primer which contained a sample index that enabled multiplexing (Table S1). First strand reverse transcription was performed for 1 hour at 42C, before being heat inactivated for 5 min at 70C, using the Smartscribe Reverse Transcriptase (Clontech). Second strand synthesis and PCR was performed with KAPA 2x master mix and an ISPCR primer for 15 cycles (37C for 30 minutes, 95C for 3 minutes, 98C for 20 seconds, 67C for 15 seconds, 72C for 8 minutes, 72C for 5 minutes, 4C hold). cDNA was cleaned up using SPRI beads at a ratio of 0.8:1. The cDNA was quantified using the qubit and subsequently pooled together. cDNA was then split for a non size-selected or a size-selected R2C2 library. To size-select, the cDNA was run on a 1% low melt agarose gel. cDNA above 2kb was excised and purified by digesting the gel with beta-Agarase and extracting the DNA from the solution using SPRI beads.

### R2C2 Library Generation

200ng of pooled cDNA libraries were circularized using 200ng of DNA splint (Table S1) using NEBuilder HiFi DNA assembly master mix (NEB E2621). Splints were designed so that their ends were complementary to the ends of the cDNA. Non-circularized DNA was digested overnight at 37 C using ExoI, ExoIII, and Lambda exonuclease (NEB) before heat inactivation for 20 min at 80 C. The reaction was cleaned using SPRI beads at a 0.85:1 (bead:sample) ratio. The cleaned, circularized library was then used for rolling circle amplification (RCA) using Phi 29 (NEB) with a random hexamer primer for 18 hours at 30 C before a heat inactivation for 10 min at 65 C. RCA product was debranched using T7 endonuclease (NEB) for 2 hr at 37 C and then cleaned and concentrated using a Zymo DNA Clean and Concentrator column (Zymo D4013). The debranched RCA product was run on a 1% low-melt agarose gel. DNA longer than 8kb was excised and extracted. The resulting R2C2 DNA was used as input into ONT library preparation.

### ONT Sequencing

ONT libraries were prepared from R2C2 DNA using the ONT ligation kit (ONT SQK-LSK110 or SQK-LSK114) and sequenced on ONT MinION and PromethION flow cells (R9.4.1 or R10.4). Flow cells were flushed with DNAse I and loaded with additional libraries 24 hours after initial loading to increase sequencing throughput.

### ONT Data Processing

Raw nanopore sequencing data were basecalled using the “sup” setting of guppy6. R2C2 full length consensus reads were generated and demultiplexed using C3POa (v2.4.0).

### Data Analysis

#### Gene-level

R2C2 data was aligned using minimap2 (Li 2018) to the GRCm39 reference genome and featureCounts was used to quantify the data set at the gene level. Gene level count data was quantile normalized using Normseq. Normalized read counts were used as input for metaCycle using the JTK_CYCLE algorithm to evaluate oscillations. Public RNA-seq reads from Atger et al. (*48*)and Zhang et al. (*22*)were aligned using STAR aligner (Dobin et al., 2013) to the GRCm39 mouse reference genome. Similar to R2C2 analysis, featureCounts was used to quantify these data sets at the gene-level and the resulting gene counts were quantile normalized using Normseq.

#### Isoform-level

Isoforms were called by the Mandalorion Isoform analysis pipeline (v4.5). Isoforms were also quantile normalized using Normseq before using metaCycle with the JTK_CYCLE algorithm to evaluate oscillations. Isoforms were classified using SQANTI3 v5.1 (Pardo-Palacios et al., 2023).

All isoforms were visualized using the GenomeBrowserShot script included in Mandalorion.

#### CDS-level

We combined read counts from all isoforms encoding the same amino acid sequence as reported by SQANTI3. CDS read counts were quantile normalized using Normseq before using metaCycle with the JTK_CYCLE algorithm.

#### TSS-level

Transcription start sites and start site counts were reported and quantified by Mandalorion. TSS read counts were quantile normalized using Normseq before using metaCycle with the JTK_CYCLE algorithm.

PolII, H3K4me3, and H3K27ac ChIP-seq reads were aligned to the GRCm39 mouse reference genome using Burrow-Wheeler Alignment (Li, 2013). ChIP-seq read alignments were analyzed for their density around TSSs for Fig. 6A and for their overlaps with TSSs for Fig. 6B.

Public TF Chip-seq data was downloaded from ChIP-Atlas in the form of a BED file using the following parameters: Assembly: M. musculus (mm10), Track type class: ChIP: TFs and others, Cell type Class: Liver, Threshold for Significance: 50, Track type: All, Cell type: All. Downloaded chip atlas data was then filtered to exclude ChIP experiments using female mice, before being converted to the GRCm39 genome using the liftover tool (UCSC Genome Browser). We then determine TSSs overlaps with the downloaded chip-atlas data. K-means clustering was performed using the scikit learn library in Python.

## DATA AVAILABILITY

All R2C2 consensus reads generated for this manuscript are available at the Sequencing Read Archive (SRA) under Bioproject PRJNA1151033. Genome Browser style images are available as zipped directories of png (https://users.soe.ucsc.edu/~vollmers/Circadian_BrowserImages_Zee.zip) and svg (https://users.soe.ucsc.edu/~vollmers/Circadian_BrowserImages_Zee_SVG.zip) files

## CODE AVAILABILITY

Mandalorion v4.5 is available at GitHub at https://github.com/christopher-vollmers/Mandalorion.

## Supplemental Material

**Supplemental Fig. S1:**
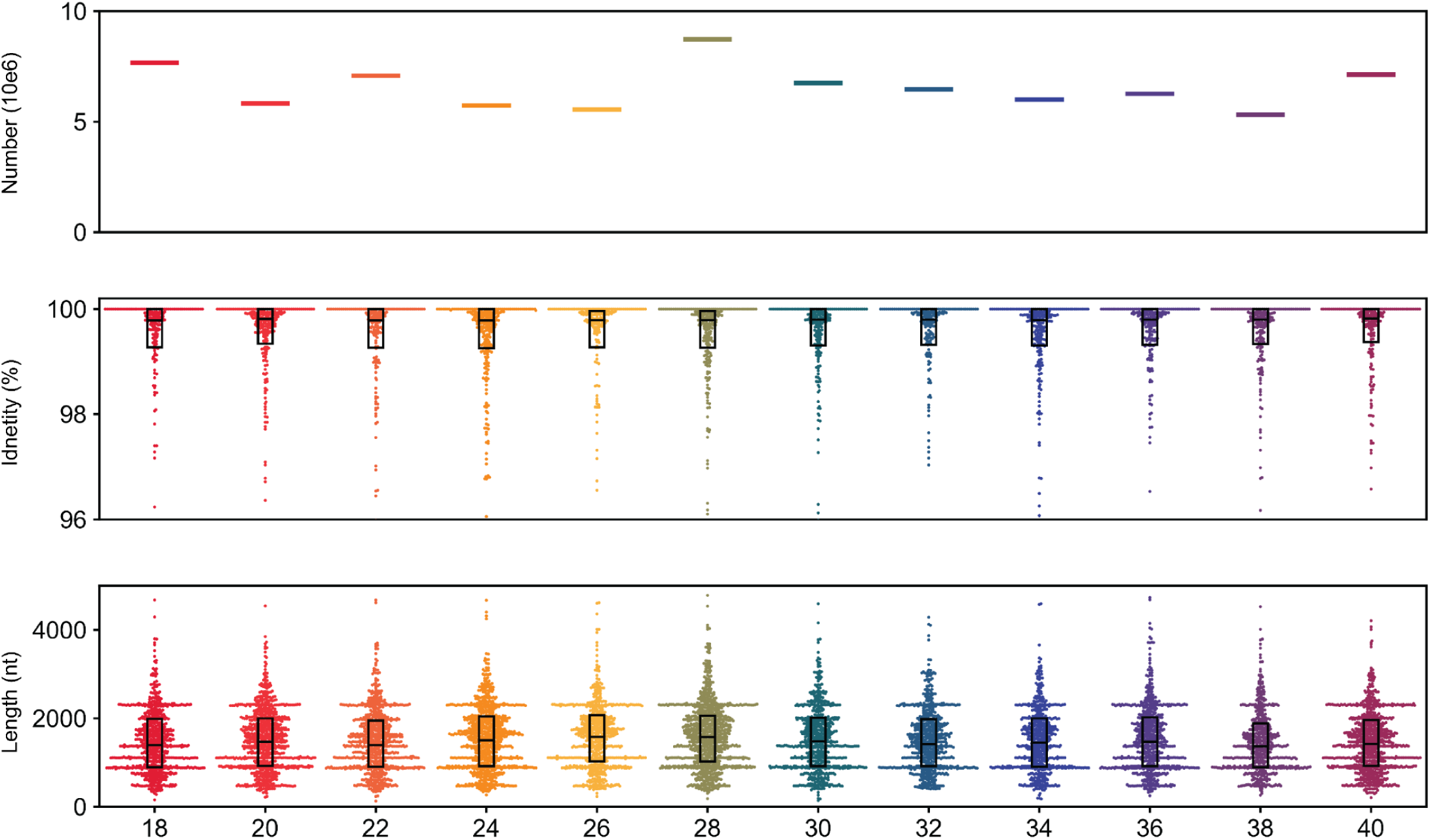
R2C2 Read Characteristics. Top panel: R2C2 consensus read counts in millions across time points. Middle panel: Swarmplot and overlaid boxplots of R2C2 consensus read identity to the mm39 genome. Bottom panel: Swarmplot and overlaid boxplots full-length cDNA molecules sequenced by R2C2.

**Supplemental Fig. S2:**
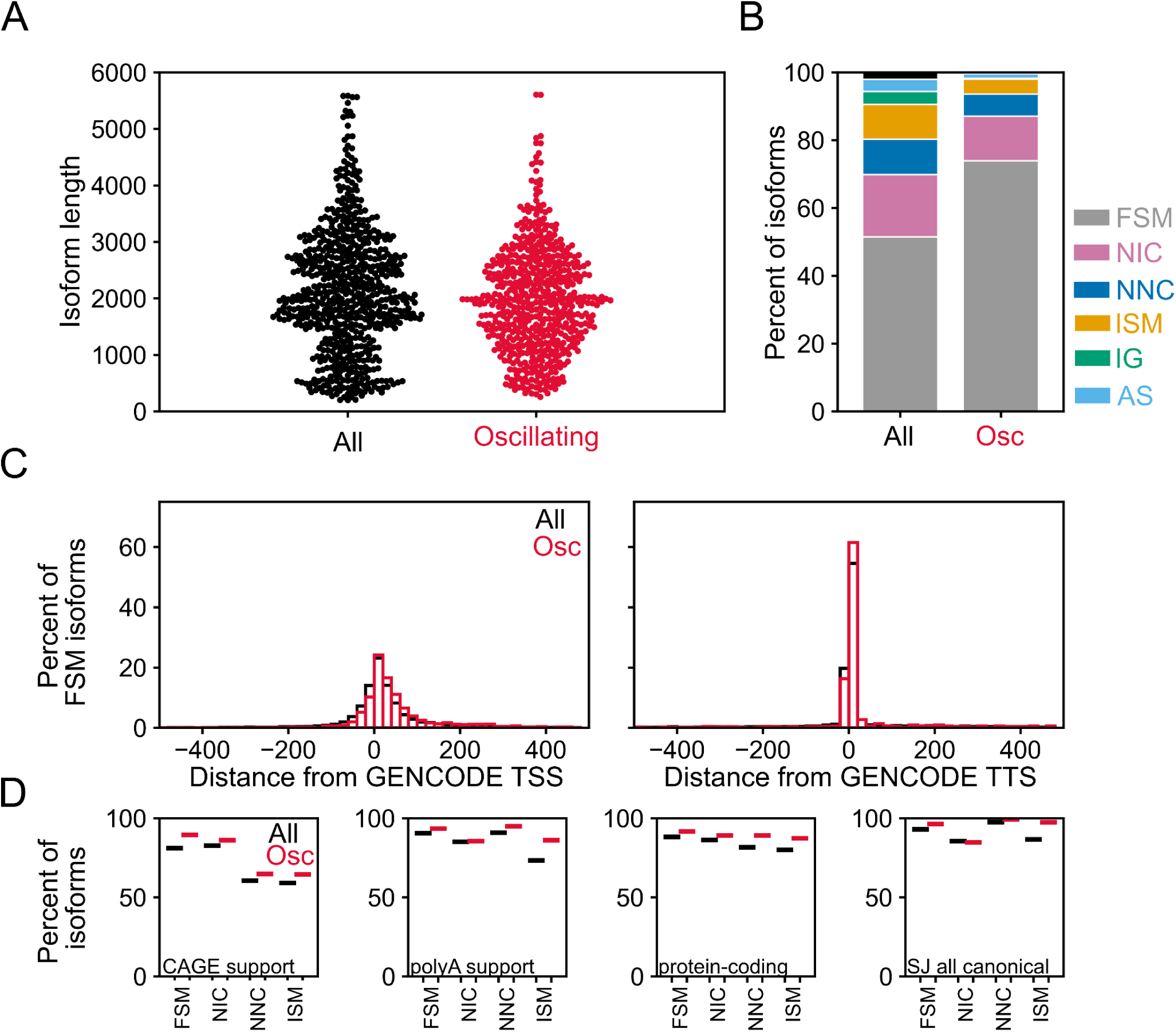
Oscillating isoform characteristics. For All and Oscillating isoforms the plots show A) Swarmplots of the length of isoforms shown as swarmplots. B) Structural isoform information determined by SQANTI shown as stacked bars. C) Histograms of distances of TSSs and TTSs of the FSM isoforms to annotated GENCODE TSSs and TTSs, respectively. D) Plots of the percentages of isoforms with characteristics determined by Mandalorion. Isoforms were split by structural category. Characteristics shown are “CAGE support”, i.e. an isoform’s TSS is close to an TSS in refTSS, “polyA support”, i.e. an isoform’s TTS is close to a polyA signal, “protein-coding”, i.e. an isoform is predicted to have a CDS, and “SJ all canonical”, i.e. all splice-junction of an isoform are canonical.

**Supplemental Figure S3:**
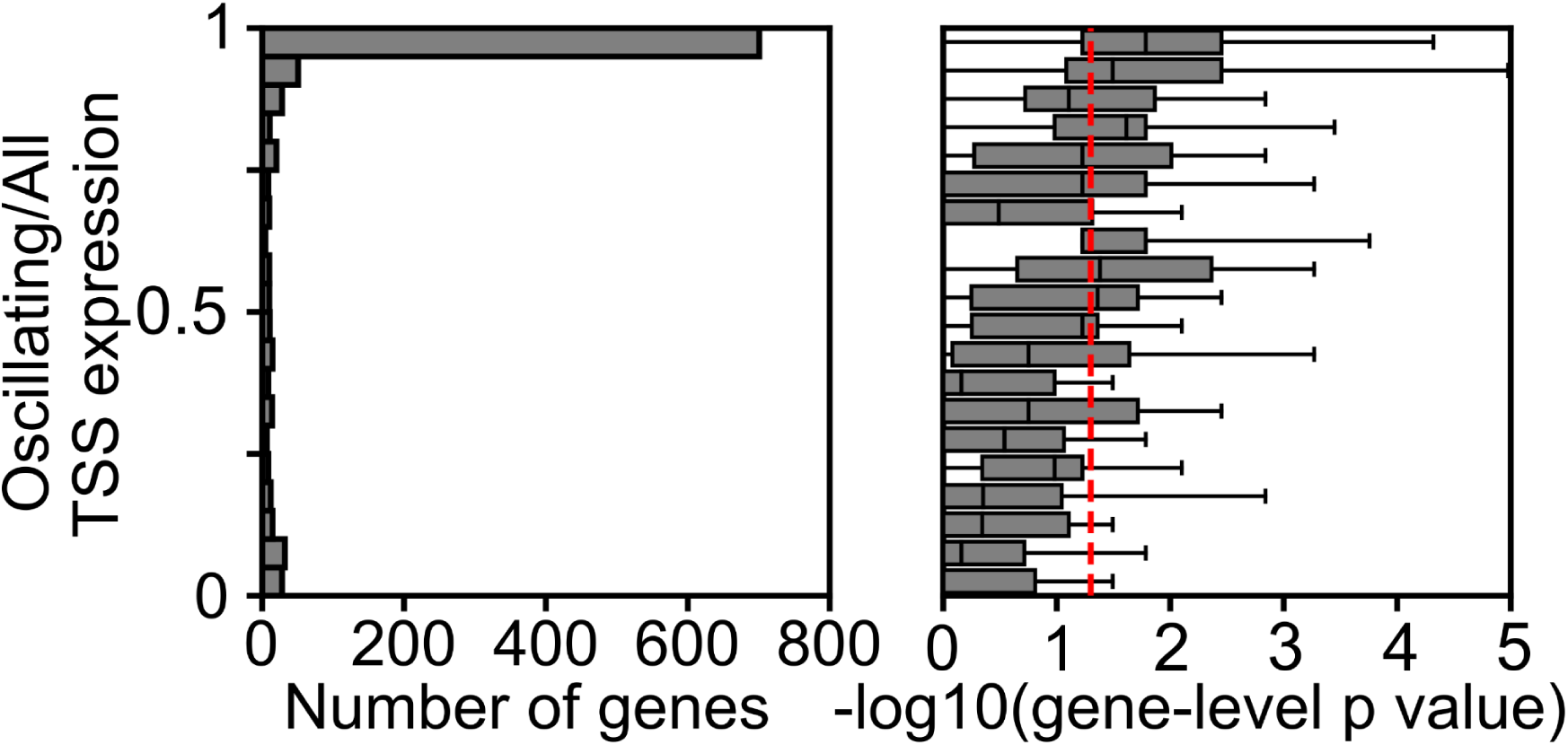
Oscillating TSSs are likely to be the dominant TSS of their gene. Left: Histogram of genes with at least one oscillating isoform binned by their expression ratio of oscillating TSS to all TSS. Right: Boxplots showing gene-level p-Value (-log10 converted) distributions of across the corresponding bins on the left. The dashed red line indicates the p-value cutoff of 0.05

**Supplemental Fig. S4:**
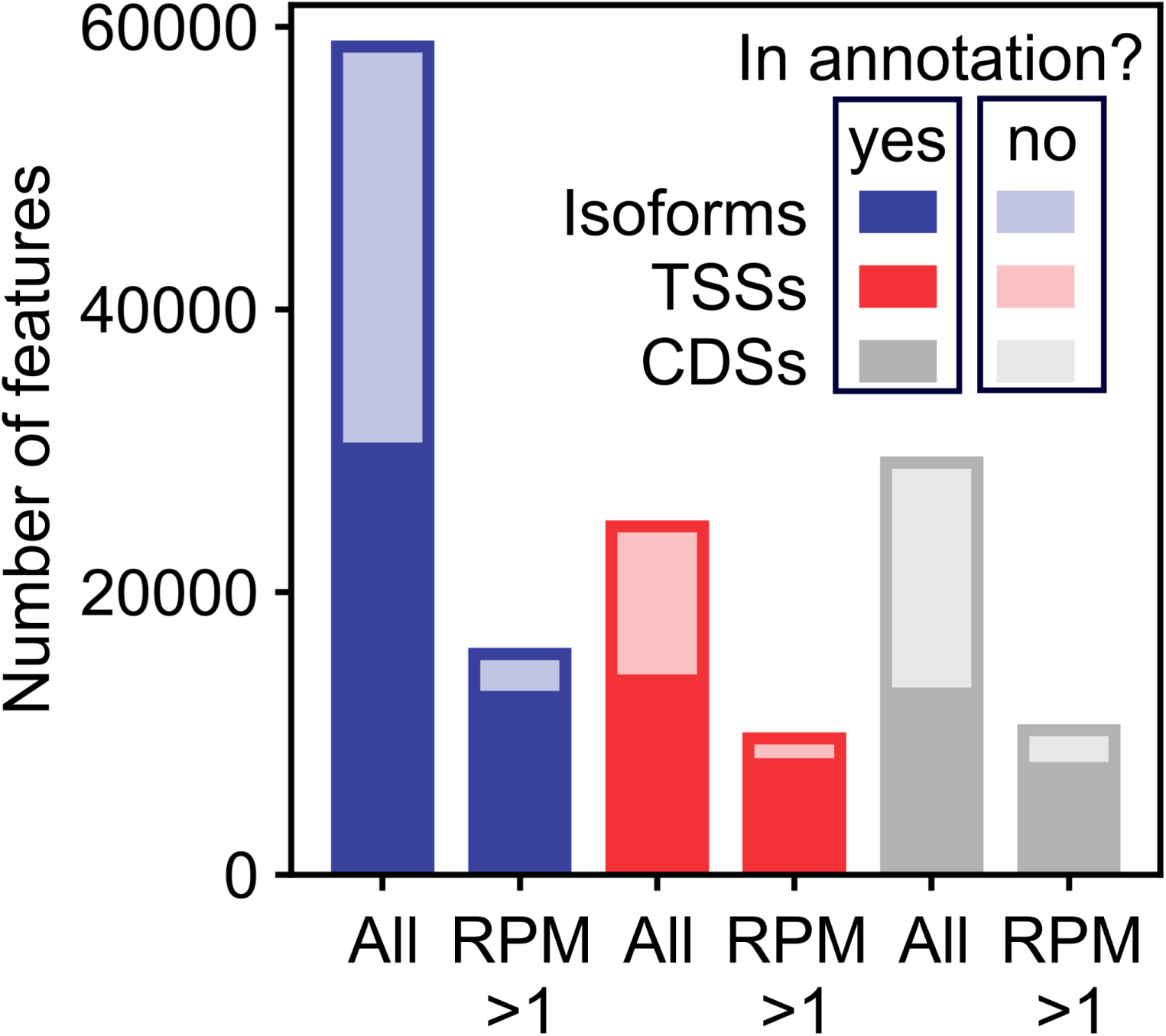
Lowly expressed features of the transcriptome arw less likely to be previously annotated. The number of Isoforms, TSSs, and CDSs detected in this study is shown as stacked bar plots. Stacked bars with different shading show features present or absent from the GENCODE vM30 annotation. Separate stacked bars show numbers of all features and those with expression over 1 RPM.

**Supplemental Table S1:**
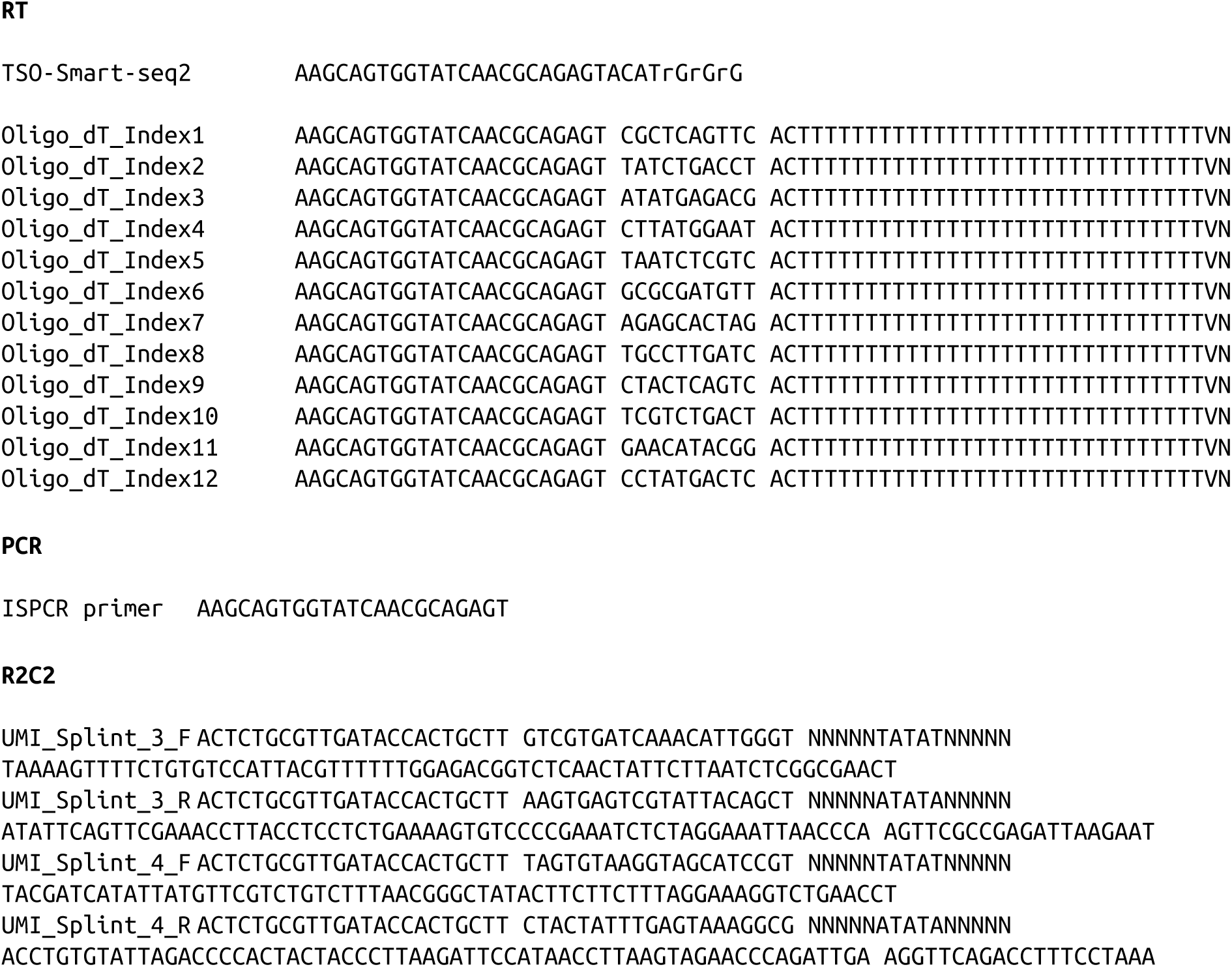
Oligos used in this study.

